# Decoding the Infant Mind: Multichannel Pattern Analysis (MCPA) using fNIRS

**DOI:** 10.1101/061234

**Authors:** Lauren L. Emberson, Benjamin D. Zinszer, Rajeev D. S. Raizada, Richard N. Aslin

## Abstract

The MRI environment restricts the types of populations and tasks that can be studied by cognitive neuroscientists (e.g., young infants, face-to-face communication). FNIRS is a neuroimaging modality that records the same physiological signal as fMRI but without the constraints of MRI, and with better spatial localization than EEG. However, research in the fNIRS community largely lacks the analytic sophistication of analogous fMRI work, restricting the application of this imaging technology. The current paper presents a method of multivariate pattern analysis for fNIRS that allows the authors to decode the infant mind (a key fNIRS population). Specifically, multi-channel pattern analysis (MCPA) employs a correlation-based decoding method where a group model is constructed for all infants except one; both average patterns (i.e., infant-level) and single trial patterns (i.e., trial-level) of activation are decoded. Between subjects decoding is a particularly difficult task, because each infant has their own somewhat idiosyncratic patterns of neural activation. The fact that our method succeeds at across-subject decoding demonstrates the presence of group-level multi-channel regularities across infants. The code for implementing these analyses has been made readily available online to facilitate the quick adoption of this method to advance the methodological tools available to the fNIRS researcher.

## Introduction

The goal of cognitive neuroscience is to use the relationship between activity in the brain and cognitive operations to understand how the mind works. In the last two decades, the use of fMRI has vastly expanded our window on the neural correlates of human cognition. Initially, fMRI analyses predominantly facilitated brain mapping: Experiments could tell us where in the brain clusters of voxels show differential BOLD signals to two or more stimulus conditions. With the addition of multivariate analysis techniques (e.g., multivoxel pattern analysis, MVPA), more sophisticated questions can be asked, such as whether the *pattern* of BOLD can discriminate between two or more stimulus conditions. Multivariate analyses are an important advance as they have shifted the focus of cognitive neuroscience from mean activation differences to the information contained within patterns of brain activity (see Raizada, Tsao, Liu, & Kuhl, 2010 for example).

MVPA studies have provided compelling evidence that the adult brain contains distributed patterns of neural activity (Haynes, 2015; Norman, Polyn, Detre, & Haxby, 2006; Serences & Saproo, 2012). Studies have demonstrated the power of MVPA to decode natural visual images (Kay, Naselaris, Prenger, & Gallant, 2008), noun identity (Mitchell et al., 2008), and speaker identity (Formisano, Martino, Bonte, & Goebel, 2008). Indeed, the use of multivariate methods has become widespread, with thousands of papers employing this method with fMRI data. Analogous multivariate methods having also been developed for EEG (e.g., Simanova, van Gerven, Oostenveld, & Hagoort, 2010), MEG (e.g., Cichy, Pantazis, & Oliva, 2014), and intracranial recordings (e.g., Liu, Agam, Madsen, & Kreiman, 2009).

However, despite fMRI’s power to address foundational questions in cognitive neuroscience, it is not universally applicable to all kinds of subject populations. For example, early developmental populations (e.g., infants) have not yet been successfully scanned while awake. Many clinical populations such as those with acute anxiety disorders, movement disorders, or cochlear implants also cannot participate in MRI experiments. Furthermore, the scanner environment greatly restricts the types of cognitive tasks and abilities that can be investigated. Cognitive abilities, such as face-to-face communication or motor movements, cannot be assessed while laying supine and motionless inside the tightly enclosed and noisy bore of the scanner. Thus, the MRI environment drastically restricts the kinds of tasks and populations that can be studied.

Functional near-infrared spectroscopy (fNIRS) is a complementary neuroimaging modality that has gained popularity in recent years for its ability to deal with many of these constraints that limit the use of fMRI. The fNIRS device records the same physiological substrate as fMRI (i.e., changes of blood oxygenation in the cortex arising from neurovascular coupling) using scattered infrared light instead of magnetic fields and radio waves. Specifically, fNIRS requires participants to wear a cap embedded with detectors and emitters of near-infrared light, similar to a pulse oximeter. A detector-emitter pair forms a *NIRS channel* within which cortical hemodynamic responses can be recorded (Figure 1C). Crucially, this method for recording the hemodynamic response is free from requirements that make the MRI environment so restrictive. No magnetic fields are needed, and the device is not sensitive to local electrical interference. Thus, it is safe for participants with metal or magnetically sensitive implants, and the machine is highly portable to many types of environments. The fNIRS device is silent, the cap is relatively comfortable to wear, and the measurements are more robust to movement than MRI. A different imaging modality, EEG, also has the advantages of being quiet and using sensors that can be attached directly to the head. However, fNIRS has significantly better ability to spatially localize neural signals than does EEG, which has poor localization due to the conduction of the electrophysiological signals throughout the head. In other words, although the coverage of a single fNIRS channel is large, one can be highly confident that the signal is specific to that region.

While there are clear limitations to fNIRS (e.g., the absence of an anatomical image, lower spatial resolution, and the ability to record only from the surface of the cortex), it provides an important tool to expand cognitive neuroscience into areas of inquiry previously beyond our methodological reach (see recent reviews from a developmental perspective: Aslin, Shukla, & Emberson, 2015; Gervain et al., 2011; Lloyd-Fox, Blasi, & Elwell, 2010).

A major hurdle to the widespread acceptance of fNIRS, as an alternative to fMRI, fNIRS data analysis techniques have historically lagged behind fMRI in sophistication. Part of this analytic immaturity arises from inherent methodological limitation of fNIRS. For example, NIRS systems do not collect the anatomical images that enable fMRI signals to be spatially localized (see Sarah Lloyd-Fox et al., 2014 for recent advances in this area). However, other aspects of fNIRS data analysis have fallen behind for reasons not inherent to the method. For example,only recently have fNIRS researchers employed corrections for multiple comparisons (see Tak & Ye, 2014 for a review of the history of statistical techniques with fNIRS) or defined prior neuroanatomical regions of interest validated by structural MRI templates (Emberson, Richards, & Aslin, 2015; Zinszer, Chen, Wu, Shu, & Li, 2014). While fNIRS methodology has greatly progressed, this neuroimaging modality will only reach its potential as a complementary neuroimaging technique if it achieves a high level of analytical sophistication.

**Figure 1.**
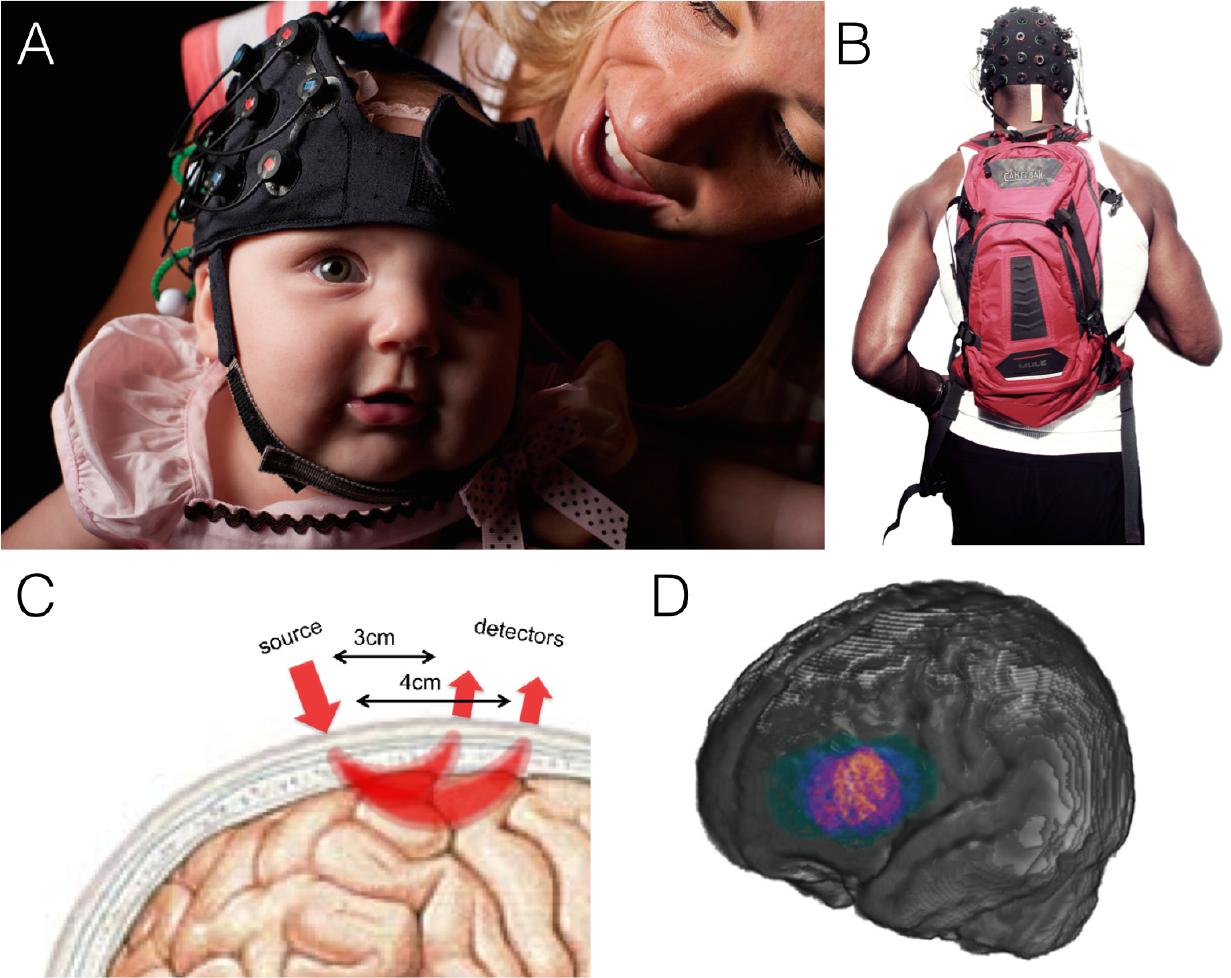
Functional Near-Infrared Spectroscopy (fNIRS) records cortical hemodynamic responses in populations that cannot comfortably be inside the MR scanner such as (A) young infants (photo credit: J. Adam Fenster). Moreover, (B) fNIRS recordings can happen “in the wild” to have more naturalistic cognitive tasks as illustrated by the NIRx Sport system. (C) Pairs of detectors and emitters form an fNIRS channel (from Gervain et al., 2011 with permission) which covers a localizable region of the cortex, e.g., (D) the spatial extent of a single fNIRS channel over a population of 6-month-old infants (from Aslin et al., 2015 with permission).

In this paper, we propose an important step forward for fNIRS data analysis by presenting a simple, easily implemented method for conducting multivariate pattern analyses with fNIRS data gathered from infants. Referencing the analogous method in fMRI, we refer to this method as **Multichannel Pattern Analysis** or **MCPA**. MCPA is computationally fast and immediately applicable to a variety of preprocessing routines already used in the fNIRS field. Moreover, we’ve made the code to implement these analyses freely available, along with the data sets used in the current paper^1^ The code can be used as a modular function with a variety of other data sets or can be readily adapted to future applications.

The application of MCPA to fNIRS data has the potential to extract substantially greater detail about the neural correlates of cognition than can be revealed using univariate statistics. Indeed, prominent results in the fMRI literature have found that hemodynamic responses encode significant information about participants’ cognitive states without producing a robust univariate contrast (e.g., Raizada, Tsao, Liu, & Kuhl, 2010; Kok, Failing, & de Lange, 2014). In the present study, we demonstrate this same sort of result for the first time using fNIRS in infants: as is described in more detail below, we show that when the classifier’s task is to distinguish between two conditions which are both audio-visual but which differ in the specific nature of the audiovisual stimuli (faces-and-music, versus fireworks-and-speech) our multichannel decoding approach is able to succeed whereas purely univariate analysis fails.

However, considering multiple channels at once yields more important benefits than simply being more sensitive than univariate analyses. A more interesting distinction is the fact that although univariate activation intensity can only go up or down, multivariate patterns have similarity relations to each other, and therefore induce a structured similarity space (see Anderson, Zinszer, & Raizada, 2015 for a recent examination of similarity-based methods). Similarity measures can, for example, quantify how much a new observation matches previous observations and thus be used to classify the new observation. These sorts of questions of representational structure simply do not arise in a purely univariate framework.

Despite the success of multivariate methods for neural decoding in other imaging modalities, there are significant challenges to applying these methods to fNIRS. First, while fNIRS records local and specific hemodynamic changes in the cortex (like fMRI), the number of recording sites is typically very few (e.g., 10 to 25 channels). Second, the spatial resolution is greatly decreased relative to an fMRI voxel: A typical channel measures an approximately 2 cm^2^ region on the surface of the cortex. The spatial distribution of these recording sites across the scalp is also usually sparse, with a 2 cm separation between adjacent channels, and signals are not sampled any deeper than 2 cm into the cortical tissue. Thus, on the one hand, the number of fNIRS recording sites is similar to EEG (on the order of dozens). On the other hand, fNIRS records a spatially localized signal, unlike EEG. Although it may be difficult to conceive of successful MVPA decoding with only 10-25 “super” voxels sparsely distributed over the cortex, these are the most typical conditions under which multivariate methods are likely to be applied to fNIRS data.

The amount of data gathered within- and across-participants also significantly differs between fNIRS and other imaging modalities. Many populations that can be readily studied with fNIRS but not fMRI (e.g., infants) cannot participate for the large number of trials typically seen in MVPA designs. These fNIRS datasets typically have a greater number of participants than the average number of trials per participant. Consequently, while many fMRI MVPA studies focus on within-subject decoding techniques and avoid the problems associated with across-subject generalization, it is unclear whether decoding will be successful with the small number of trials and the high-degree of between-subject variability typical of fNIRS datasets. Thus, while multivariate methods have achieved broad success within the fMRI community (and increasingly in high-density EEG and MEG), it is unclear whether this approach will be successful given the constraints of the typical infant fNIRS dataset.

While there have been a few successful attempts applying multivariate methods to fNIRS data from adults or children, no previous study has attempted the far more challenging task of decoding fNIRS data from infants. This is likely because infant datasets always contain fewer stimulus-presentation trials and more noise than the data that can be obtained from older and hence more cooperative participants. The present study employs some similar approaches used in this prior body of work, but they also differ in crucial ways. Previous studies have decoded emotional state (Heger, Mutter, Herff, Putze, & Schultz, 2013), subjective preference (Hadi et al., 2011; Luu & Chau, 2009), item price (Misawa, Shimokawa, & Hirobayashi, 2014), vigilance (Bogler, Mehnert, Steinbrink, & Haynes, 2014), and ADHD and autism diagnoses (Ichikawa et al., 2014) from fNIRS recordings. While the results are promising (decoding accuracy between 60-70%), these initial attempts fall short of providing a methodological foundation from which the fNIRS community can build further multivariate analyses. First, these studies have all employed Support Vector Machine (SVM) algorithms for classification. SVMs are a machine learning technique that seek a set of weights for the hemodynamic signal from each voxel or channel that best predicts classification accuracy with as large a margin as possible between the classes to be distinguished (Norman et al., 2006). However, typical of many machine learning techniques, SVMs typically require data from a large number of stimulus-presentation trials from each participant in order to converge, and the weights of a successful classifier are difficult for researchers to interpret in relation to their original research questions, making it problematic for wider adoption by the fNIRS community.

Second, these previous studies typically assessed adults rather than the infant and clinical populations that are uniquely suited for fNIRS imaging. The current limitation of multivariate fNIRS methods to typically functioning adults has thus not inspired broader application. Many populations typically studied in fNIRS cannot contribute data a large number of trials in an experiment and are more difficult to test. Thus, these studies (with the exception of Ichikawa et al., 2014) have larger pools of data from each individual participant available for the classification (typically 50+ samples per participant) but these large within-subject datasets would be extremely difficult to achieve in infants or many clinical populations.^2^ Given the great deal of neural variability across individuals, especially in clinical populations and early in development, performing decoding across individuals is quite difficult and has typically been more challenging for MVPA studies using fMRI. However, as we will demonstrate, across-infant multivariate inferences can produce highly accurate results while overcoming one of the most common practical limitations of infant research (limited numbers of trials per infant).

The current study extends previous work in three major respects:

1. We employ a correlation-based decoding method that is computationally simple, fast, and easily interpretable. The code for implementing all the analyses and a sample dataset are included to encourage immediate application and broad adoption in the fNIRS community.
2. The present study is the only reported multivariate analysis with infants. A large number of researchers employing fNIRS are doing so to probe the neural correlates of early development, but they have exclusively used univariate methods (Aslin et al., 2015). It is currently unknown whether decoding the infant brain is possible using MCPA, especially given the small number of trials contributed by infants in a typical experiment.
3. The current MCPA analysis performs all classifications *across* infants. Data from an infant who does not contribute to the group model are classified by aggregating data across other infants in the dataset. The between-subjects approach takes advantage of the larger number of available participants in an experiment than trials completed by each participant.

Our overall goal is to determine whether multichannel pattern analysis (MCPA) can decode the infant brain. Specifically, we examine whether patterns of activation across multiple NIRS channels can accurately predict the patterns of activation in an infant who did not participate in the creation of a *group model*. We examine the decoding accuracy for each *test infant’*s activation patterns as that infant’s data are iteratively removed from the group model. We refer to the resulting classification of each test infant’s condition averages as *infant-level* decoding. We also examine the much more difficult task of classifying each of the test infant’s individual trials (*trial-level* decoding) based only on the group model to which the left out test infant did not contribute.

In sum, we ask whether a group model of neural responses created from several infants sufficiently generalizes to classify observations from a new infant (the test infant). Our measure of success is whether decoding produces accurate predictions (i.e., significantly above chance) when these MCPA methods are iterated across the entire sample of infants. To determine whether this method is robust and generalizable to new experiments, we will also apply it across two separate groups of infants in different experiments presenting different levels of decoding difficulty.

## Methods

### Multichannel Pattern Analysis

Multichannel pattern analysis (hereafter, MCPA) uses a group model, derived from multichannel patterns of activation, to decode unlabeled patterns of activation that did not contribute to the group model. We begin by describing how ***multichannel patterns*** are calculated, then how the ***group model*** is created, and finally how this group model is used to decode ***unlabeled*** multichannel patterns. All analyses were conducted in MATLAB (version R2013a, 8.1.0.604) with custom analysis scripts.

First, multichannel patterns are constructed by including information for all channels, for each trial, for every infant. MCPA treats each channel as an independent observation. To derive a multichannel pattern, oxygenated hemoglobin (HbO) is averaged over an *a priori* determined time-window (i.e., up to *t* measures after stimulus onset) for each channel (*chan*). This is described in Equation 1 and Figure 2A.

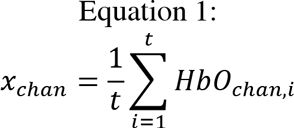

The average oxygenated response (*x_chan_*) is determined for each channel for the same time-window and then combined with the corresponding averages from the other channels into a single vector to create a *multichannel pattern* 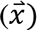 for a single time-window (e.g., single trial), for a single infant (see Figure 2A). These multichannel patterns are vectors of dimension *n* where *n* equals the number of channels.

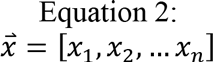

Multichannel patterns can then be averaged across all the trials (*r*) of a given stimulus condition within infants to give rise to an ***infant-level multichannel pattern*** for that stimulus condition (i.e., in an experiment with two conditions, two such vectors are calculated, one for each condition).

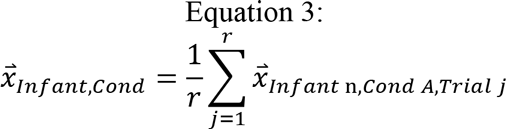

To construct the ***group model***, infant-level multichannel patterns, for each stimulus condition, are averaged to produce a group-level multichannel pattern 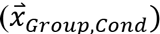. However, it is important to note that not all infants are included in this average: MCPA employs a leave-one-out method so a single ***test infant*** is removed from the average and the group model is created from N-1 infants for the current experiment. See Figure 2B. This leave-one-out process is iterated N times so that each infant has been left out and will serve as the test infant, and thus, the group average has been recomputed N times from the remaining (non-test) infants.

The multichannel patterns are compared to one another using Pearson correlation. Because the multichannel pattern vectors have the same number of dimensions (the number of channels) at all levels of analysis, we can compare a group-level model to (1) the infant-level multichannel pattern for the test infant or (2) the multichannel patterns for single trials from the test infant. We will refer to the former as ***infant-level decoding*** and the latter as ***trial-level decoding***. While these two types of decoding are derived from the same basic principles, they differ in that the infant-level decoding has multiple infant-level multichannel patterns to decode simultaneously (the number is equivalent to the number of stimulus conditions, but in this paper, we compare across two conditions exclusively), whereas trial-level decoding, by definition, has only a single multichannel pattern to decode. These differences methodology between the two types of decoding result in subtle differences in how the group model is used to decode these multichannel patterns of activation. We now describe these differences in detail.

Infant-level decoding uses the group model to predict the infant-level multichannel patterns of the test infant for each stimulus condition. Specifically, the test infant’s multichannel patterns are decoded by determining which permutation of condition labels yields the greatest overall correlation to the group-level model. For two conditions, A and B, two Pearson correlations are computed as follows:

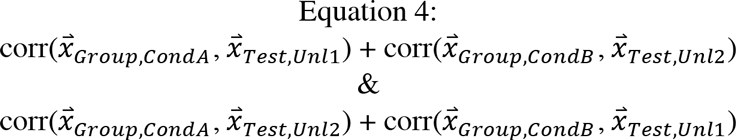

The sums of the two Pearson correlations are compared. Decoding is considered successful if the sum of the correlations with the correct labels is greater than the sum of the correlations with the incorrect labels. In other words, if Unlabeled1 is the test infant’s average response to ConditionA and Unlabeled2 is the test infant’s average response to Condition B, one could consider that whichever permutation yields the greatest sum of correlations (A-1, B-2 vs. A-2, B-1) provides the best estimate of labels for the new, unlabeled responses. If the new labels are correct, then decoding of this test infant with the group model is considered accurate, and if not, then decoding is considered inaccurate (a 1 or a 0 is assigned for this test infant, respectively).

**Figure 2.**
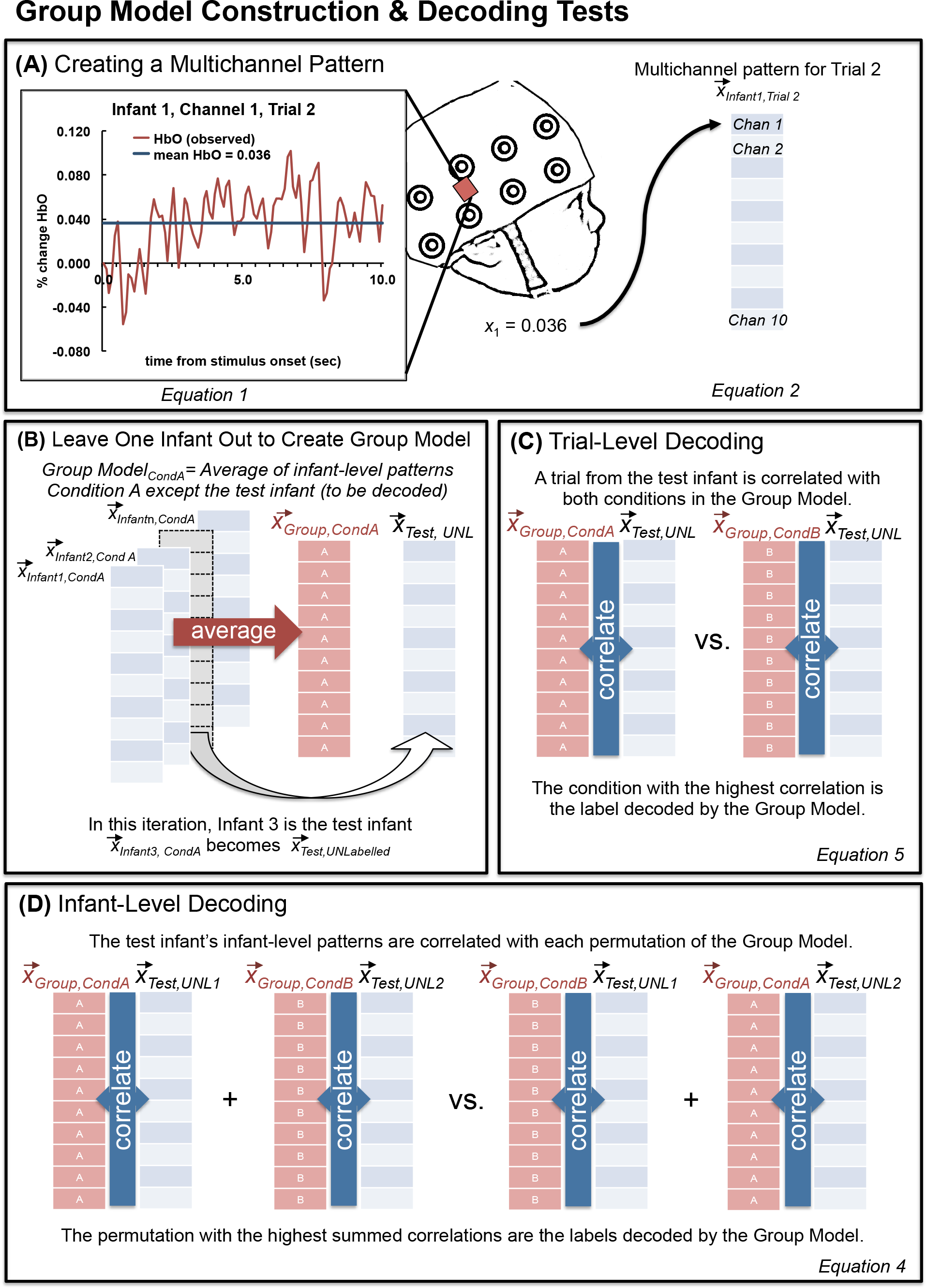
Illustration of the methods of multichannel pattern analysis (MCPA).

For trial-level decoding, the test infant’s multichannel patterns for each unlabeled trial are correlated with each condition from the group model. If the correct label yields the greater correlation then we consider decoding to be successful. For example, a correctly decoded ConditionA trial would be represented by the following equation:

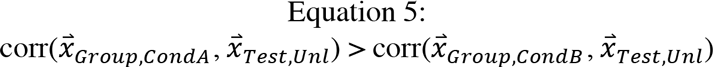

After binary decoding accuracy (1/0) is recorded for each trial for a given test infant, average accuracy for each condition is derived by averaging decoding accuracy for all trials of a given condition.

Once decoding (infant-level or trial-level) is complete for a given test infant, this procedure is iterated throughout the population of infants until each infant has been a test infant and their patterns of activation have been decoded. Specifically, after decoding is complete the test infant is reintegrated and the multichannel patterns from another infant are removed (i.e., the new test infant) before a new group level model is created. To conduct decoding for *N* infants in a given experiment, *N* group models are created with a unique model for each infant.

These methods can be applied to three or more stimulus conditions, but the present study will focus on the case of only two conditions. Extension to a greater number of conditions is further addressed in the Discussion.

### Application of MCPA Method to Infant fNIRS Data

We applied our MCPA method to two previously collected infant fNIRS datasets. Both datasets were obtained from a Hitachi ETG-4000 and a custom cap and optical fibers built especially for fNIRS data collection with infants. FNIRS data were sampled at 10Hz. Twenty-four channels were used in the NIRS cap, with 12 over the back of the head to record bilaterally from the occipital lobe, and 12 over the left side of the head to record from the left temporal lobe. The channels were organized in two 3×3 arrays, and the cap was placed so that, for the lateral array, the central optode on the most ventral row was centered over the infants’ left ear and, for the rear array, the central optode on the most ventral row was centered between the infant’s ears and over the inion. This cap position was chosen based on which NIRS channels were most likely to record from temporal and occipital cortex in infants. Due to curvature of the infant head, a number of channels did not provide consistently good optical contact across infants (the most dorsal channels for each pad). We did not consider the recordings from these channels in subsequent analyses and only considered a subset of the channels (7 for the lateral pad over the ear and 3 for the pad at the rear array).

During the experiment, the infant sat on a caretaker's lap in a darkened room and surrounded by a black curtain to reduce visual distraction and to separate the participant from the experimenter. Caretakers were instructed to refrain from influencing their infant, only providing comfort if needed. Participants watched the video until they consistently stopped looking, became fussy, or, in the case of dataset #2, all experimental blocks were watched.

Raw data were exported to MATLAB (version 2006a for PC) for subsequent analyses with HomER 1 (Hemodynamic Evoked Response NIRS data analysis GUI, version 4.0.0) for a standard preprocessing of the NIRS data. First, the “raw intensity data is normalized to provide a relative (percent) change by dividing the mean of the data” (HomER 1.0 manual). Then the data is low-pass and high-pass filtered (two separate steps) to remove noise such as heart-beat and blood pressure changes. Second, changes in optical density are calculated for each wavelength, and a PCA analysis was employed to remove motion artifacts. Finally, the modified Beer-Lambert law is used to determine the changes (delta) concentration of oxygenated and deoxygenated hemoglobin for each channel (the DOT.data.dConc output variable was used for subsequent analyses, see the HomER Users Guide for full details; Huppert, Diamond, Franceschini, & Boas, 2009).

#### Dataset 1: Differentiating Auditory and Visual Processing

This dataset came from an experiment in which infants either viewed a visual stimulus (a red smiley face appearing for 1 sec in a white box on the screen) or heard an auditory stimulus (a toy sound, either a rattle or a honk sound that played for 1 sec) in separate events in an event-related design. Including static presentation of the empty white square on either side of these unimodal stimulus events, these trials lasted 3-3.5 seconds (see Emberson et al., 2015 for detailed information about stimulus presentation) and were separated by a jittered baseline lasting between 4-9 seconds (mean 6.5 seconds). See Figure 3 (left panel) for a schematic of this task (i.e., which information we must decode from the neural signal).

Twenty-five (25) infants were recruited for this study (mean age = 5.7, *SD* = 0.61 months, 10 female, 2/25 infants were identified as Hispanic and 23 infants were identified by their parents as Caucasian and 2 were identified as mixed race, Caucasian + Asian, Caucasian + Native American + Black). Of these infants, 19 were included in the final data analyses, with 3 infants excluded due to poor optical contact (e.g., due to a large amount of dark hair) and 3 for failing to watch the video to criterion. Infants were recruited through the database of interested participants from the Rochester Baby Lab and were born no more than 3 weeks before their due date, had no major health problems or surgeries, no history of ear infections, nor known hearing or vision difficulties. Caregivers were compensated $10 for their visit and a token gift (e.g., a Baby Lab t-shirt, bib or tote bag).

**Figure 3.**
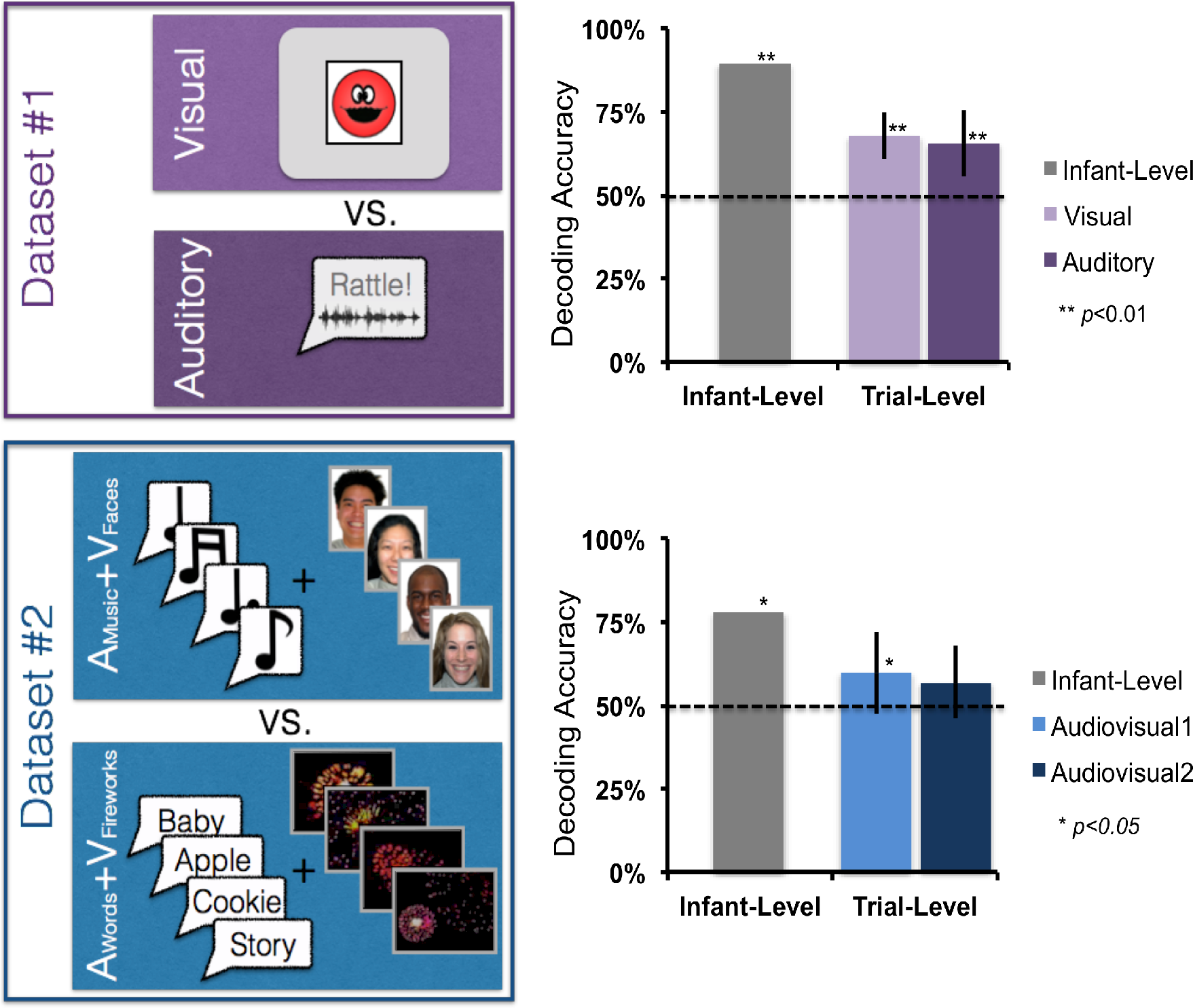
Depiction of the two datasets and the decoding results (Infant-Level and Trial-Level) for each.

Separate univariate analyses for Dataset 1 were previously reported in Emberson, Richards, and Aslin (2015). For more details, please refer to that manuscript. On average, infants watched for 6.9 trials of each of the two unimodal conditions (*SD* = 1.95). For the current MCPA analyses, the data were analyzed with a time-window of 0-10 seconds. This is a different time-window than that employed in Emberson et al. (2015), which employed a 4-11.5 second time-window. This difference reflects the fact that Emberson et al. (2015) used univariate analyses to capture the peak of the hemodynamic response function, whereas the current analysis it was important to maximize the variability of the response curve for optimal decoding. Thus, a time-window was chosen which included the rise of the hemodynamic response in the first four seconds. Parameters for selecting the optimal time window for a particular experiment are not yet clear, so while this visual inspection approach proves sufficient in the present study, the same time window may not generalize to all event-related designs. Future research will be necessary to explore the impact of time window parameters and best methods for selection.

#### Dataset 2: Differentiating Combinations of Audiovisual Processing

In this dataset, infants either viewed a block of 8 faces while listening to music (**V_Faces_+A_Music_**) or they listened to a block of 8 words while looking at dimmed fireworks (**V_Fireworks_+A_Words_**). Thus, in this dataset, successful decoding of the pattern of neural responses required distinguishing between types of audiovisual (AV) processing and not simply whether the infant is hearing something or seeing something as in Dataset #1. The audio stimuli were 8 common words familiar to infants and commonly attested in samples of infant-directed speech (apple, baby, bottle, blanket, cookie, diaper, doggie, story). The visual face stimuli were 8 smiling Caucasian female faces from the NimStim database (Tottenham et al., 2009). All stimuli had a stimulus onset interval of 1 second. The inter-stimulus interval (ISI) for visual stimuli was always ·25 seconds. The ISI for audio stimuli ranged from ·2-·3 seconds because of slight differences in word duration. The 8 stimuli in each modality were presented in shuffled order for each block. As with Dataset #1, these stimuli were followed by a jittered 4-9 second baseline (mean = 6.5 sec). See Figure 3 for a schematic of this dataset. Unrelated univariate analyses of this dataset are presented in Emberson, Cannon, Palmeri, Richards, & Aslin (under review) with more details on the experiment available in that manuscript.

Twenty-six infants were recruited based on the same criteria and using the same methods as Dataset #1. Of these, 18 infants were included in the final analysis: Infants were excluded for poor optical contact (6), not watching to criterion (1), or refusing to wear the NIRS cap (1). The remaining sample of infants had a mean age of 5.8 months (*SD*= 0.6) and consisted of 9 females and 9 males. Of the included infants, 88.9 percent heard only English at home. Two other participants heard another language from their family 60 or 90 percent of the time. Participants were identified as Caucasian (16), black (1), and Hispanic (2). Infants watched these two multimodal blocks an average of 4.9 times each (*SD* = 1.17). As with Dataset #1, the time-window investigated for MCPA is slightly different than for the corresponding univariate analyses (6-14.5 seconds — Emberson, Cannon, Palmeri, Richards, & Aslin, under review). In this block design, the rise of the hemodynamic response is less informative than the cumulative response of the eight stimuli during the exposure period. This is because the block duration is much longer than the event duration in Dataset #1 (8 sec vs. 1 sec). Thus the current analyses employ a time window of 6-12.5 seconds, which captures the period of peak activation for the stimulus block as well as the rise and fall of activation around this peak. Similar to Dataset #1, this time window also curtails the end the hemodynamic response period. Exploratory testing found that, for this particular dataset, including the beginning (e.g., starting from 0 seconds after stimulus onset), contributed a great deal of noise to the decoding, likely because the long rise time was over-represented in the averaged HbO data for the measurement window and thus did not capture sufficient variability in the responses across channels.^3^

## Results

Each individual infant’s data was decoded using a leave-one-infant-out cross validation method (also known as *n*-fold cross validation). A group model is built on all infants’ data except the *test infant*. Then this group model is used to decode the test infant’s data according to the statistical procedures described in Methods (see Figure 2 and relevant section in Methods). For each test infant, two tests were performed: (1) labeling the averaged stimulus conditions or the *infant-level* activation patterns and (2) labeling each individual trial or the *trial-level* activation patterns. The procedure is iterated through the dataset so that each of *N* infants becomes the test infant based on the decoding predictions of the remaining (*N*-1) infants, and their patterns of activation (infant-level or trial-level) are decoded based on the group average. The resulting decoding accuracy is averaged across all test infants to produce an overall decoding accuracy score: Infant-level decoding accuracy is expressed as the percentage of infants where the averaged multichannel patterns for the two conditions are correctly labeled (greater correlation for correct condition labels than the incorrect condition labels, see Equation 4). Trial-level decoding accuracy is expressed as the average percent of correctly labeled trials across infants, where correct refers to cases where the correlation is higher for the correct condition label than the incorrect condition label (see Equation 5). We apply these tests to two datasets and ask whether these decoding methods reliably predict, on an infant-level or trial-level, the obtained infant fNIRS data.

### Infant-level decoding

In Dataset #1, the infant-level multichannel patterns (averaged for each condition, Figure 2) were correctly labeled for 17 out of 19 infants (accuracy = 89%; see top panel of Figure 3, “Infant-level” data). In Dataset #2, the infant-level multichannel patterns for the two types of audiovisual trials were correctly labeled for 14 out of 18 infants (accuracy = 78%; see Figure 3, bottom panel). Because the cross validation procedure introduces dependencies between each infant that is tested against the group model, a parametric binomial test is not appropriate for testing the significance of these results. Instead, we performed a permutation-based test to create an empirical null distribution and determine the *p*-value for the observed decoding results. This procedure is described and illustrated in detail in the supplementary materials (see also Nichols & Holmes, 2003). Decoding accuracy was statistically significant in both Dataset #1 (*p*=0.001) and Dataset #2 (*p*=0.033).

Thus, we find that the group model can successfully decode two stimulus conditions in an infant’s fNIRS data with high accuracy, determining whether a test infant is either seeing or hearing a stimulus (Dataset #1), and in the much more difficult contrast between different combinations of audio-visual stimulation (Dataset #2) where successful decoding cannot rely on stimulus modality exclusively. In Dataset #2, accurate labeling requires the correct discrimination of one audiovisual combination from another, such that only subtle patterns of cortical activation in the same or overlapping regions will distinguish these two conditions rather than broad spatial differences among channels (e.g., anterior-vs.-posterior or right-vs.-left), which we test in the univariate analyses presented later in this section.

### Trial-Level decoding

For Dataset #1, labeling each trial independently for each test infant yielded 68% accuracy for Visual trials and 66% for Auditory trials (Figure 3, top panel). Each of these accuracy scores significantly exceeded chance (50%, see supplementary materials for the details of generating the null-hypothesis distribution and *p*-values; Visual: *p*<0.001; Auditory: *p*<0.001).

For Dataset #2, labeling each trial independently yielded 60% accuracy for Audiovisual-1 trials and 57% for Audiovisual-2 trials (Figure 3, bottom panel). While numerically above 50%, these accuracy scores did not robustly exceed chance (50%, Audiovisual-1: *p*=0.049; Audiovisual-2: *p*=0.128).

Thus, we find that the group model can successfully decode single trials for an infant who did not contribute to the group model (i.e., test infant). In the case of Dataset #1, both conditions were decoded significantly beyond chance. While for Dataset #2, one of the conditions was decoded beyond chance performance, the other was not suggesting some asymmetry in the information present in the fNIRS recordings.

### MCPA with Subsets of Channels

Since the current MCPA method works on multichannel patterns (i.e., vectors) of arbitrary (but consistent) length, it is also possible to perform the infant-level and trial-level analyses on subsets of the fNIRS channels, rather than using all available fNIRS channels as we reported in the previous sub-sections (10 channels).

Canonical MVPA analyses of fMRI data use voxel-stability and “searchlight” techniques to determine which voxels contain sufficient information to decode previously unlabeled trials (see Kriegeskorte et al., 2006; Mitchell et al., 2008 supplementary materials). Due to the small number of fNIRS channels and the large spatial extent of cortex over which each channel 2 (~samples cm of infant cortex, see Aslin, Shukla & Emberson, 2015), fNIRS recordings from the two datasets are not conductive to a spatial searchlight analysis. However, the small number of channels permits an exhaustive combinatorial analysis of decoding accuracy for all possible channel subsets.

This subset analysis allows us to determine which channels make the greatest overall contribution to MCPA decoding accuracy. Specifically, we consider three interrelated issues:

1. Are some channels more informative than others? With smaller subset sizes, we test the varying contribution of each channel to decoding accuracy.
2. How many channels should be included in a subset for maximum decoding accuracy? Does the inclusion of more channels result in better decoding?
3. Will the selective inclusion of highly informative channels identified in #1 (i.e., eliminating noisy channels) improve decoding accuracy?

These issues together allow us to tackle the question of whether this multivariate method can detect **spatial-specificity** in decoding accuracy. Foreshadowing our results, we will provide evidence that we can detect varying levels of decoding accuracy across channels and that these inter-channel differences can be exploited to significantly boost decoding accuracy for smaller subset sizes (points 1 and 3 respectively). Together, these results provide evidence that spatially specific decoding results can be obtained through MCPA.

In this subset analysis, we varied the number of channels included in the decoding analysis (*subset size*) from two channels to ten channels (i.e., the complete array, as reported in the foregoing results). For every subset size, _10_C_n_ possible combinations of *n* channels exist. For example, for a subset size of 2 channels, subsets would be channels 1 & 2,1 & 3, 1 & 4, etc. We tested all subsets for infant-level decoding.

### Are Some Channels More Informative Than Others?

To test whether some channels are more informative than others, subset sizes of 2 to 10 channels were compared for infant-level decoding accuracy, with all possible combinations of channels tested in _10_C_n_ subsets. For each subset, the average decoding accuracy was assigned to each channel. Once all subsets are decoded, each individual channel’s accuracy is defined as the average accuracy for all the subsets it participated in.

Figure 4 presents average infant-level decoding accuracy for each of the 10 channels across subset sizes from 2 to 10 for both datasets. Three patterns are evident. First, as predicted, mean decoding accuracy across channels declines with decreasing subset size (i.e., decreasing numbers of channels participating in the decoding algorithm). Further, subtle differences between channels become visually apparent as subset size decreases from 10 to 5 and more pronounced with subset sizes of 2 and 3. Interestingly, we find that although both Datasets involve stimulus conditions that contain auditory and visual information, there is some difference in which channels appear to be the most informative: for Dataset #1, the three most informative channels (numerically) are 1, 3 and 8; for Dataset #2, the three most informative channels are 1, 2, and 9. Information about the spatial location of these channels can be found in the Supplementary Materials but generally, channels 1 through 3 are located in the occipital lobe spanning from V1 to LOC, the remaining channels are distributed through the temporal cortex (5 channels) and the prefrontal cortex (2 channels). Thus, we find that as subset size decreases, there is more diversity in mean decoding accuracy across channels, with prominent differences revealed between channels in subset sizes 2 and 3.^4^ In the following subsections, we estimate a statistical model to test the effects of set size and the differences between channels and investigate whether these differences can inform the selection of channels for better decoding accuracy.

**Figure 4.**
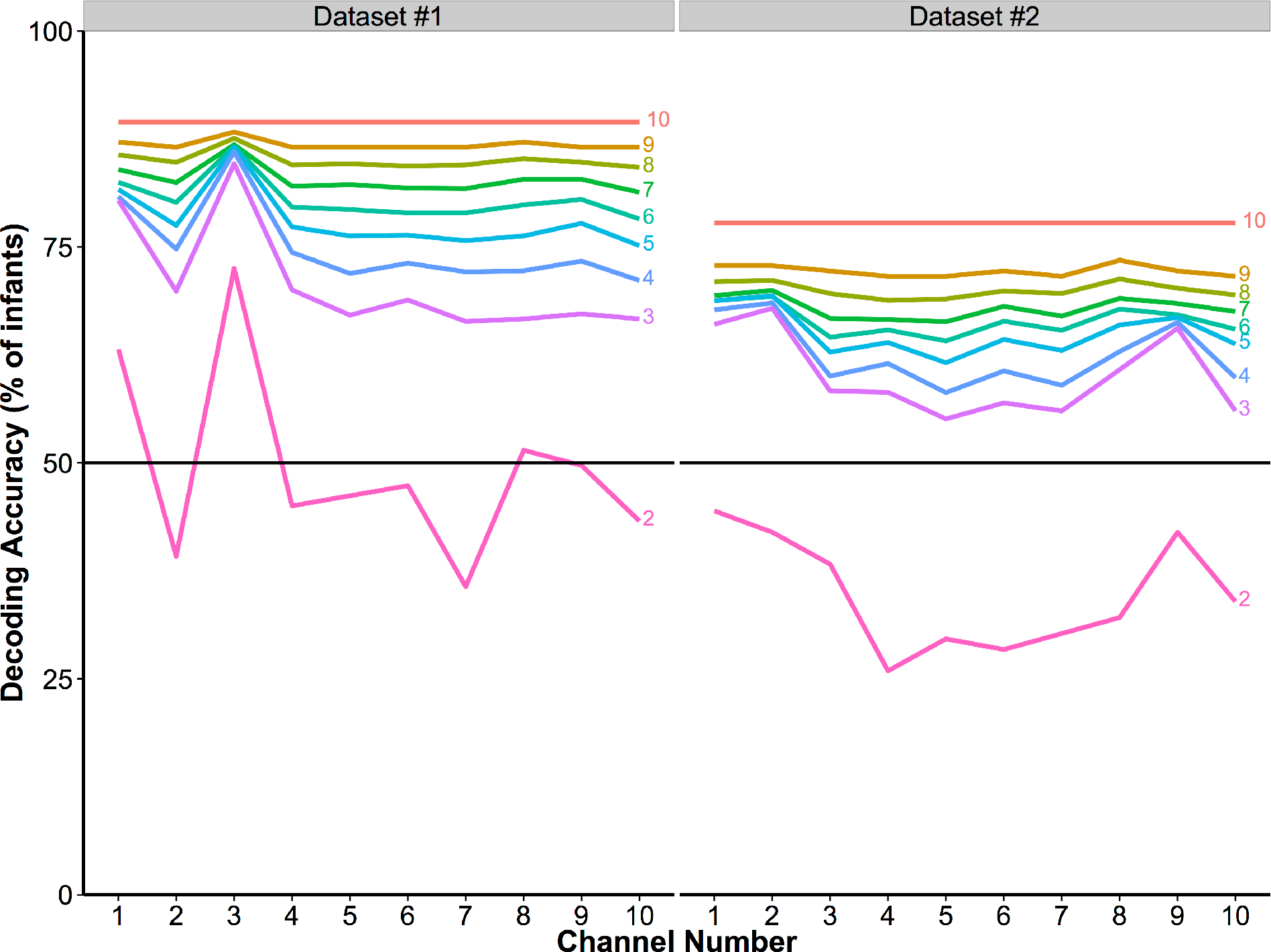
Accuracy for each of the 10 NIRS Channels for Dataset #1 (left) and Dataset #2 (right) in different subset sizes (from 2 to 10 channels with each line labeled at the right with the subset size).

Another pattern evident in Figure 4 is that the difference in accuracy between Datasets #1 and #2, when all 10 channels are used (Figure 3, as reported above), is also present across all subset sizes. We turn to a detailed investigation of differences in accuracy across subset sizes in the next subsection.

### Are There Differences in Decoding Accuracy with Subset Size?

To determine how decoding accuracy changes with subset size, accuracy over all subsets was determined for each subset size and for both datasets. As shown in Figure 5, the difference in average decoding accuracy increases with larger subset sizes and peaks for both datasets when the subset size equals the total number of channels (subset size = 10). Thus, even though all channels are included across subsets, when subset sizes are less than the total number of channels (< 10), the maximum average decoding accuracy (across all subsets of the same size) improves as the number of channels simultaneously considered during decoding increases.

**Figure 5.**
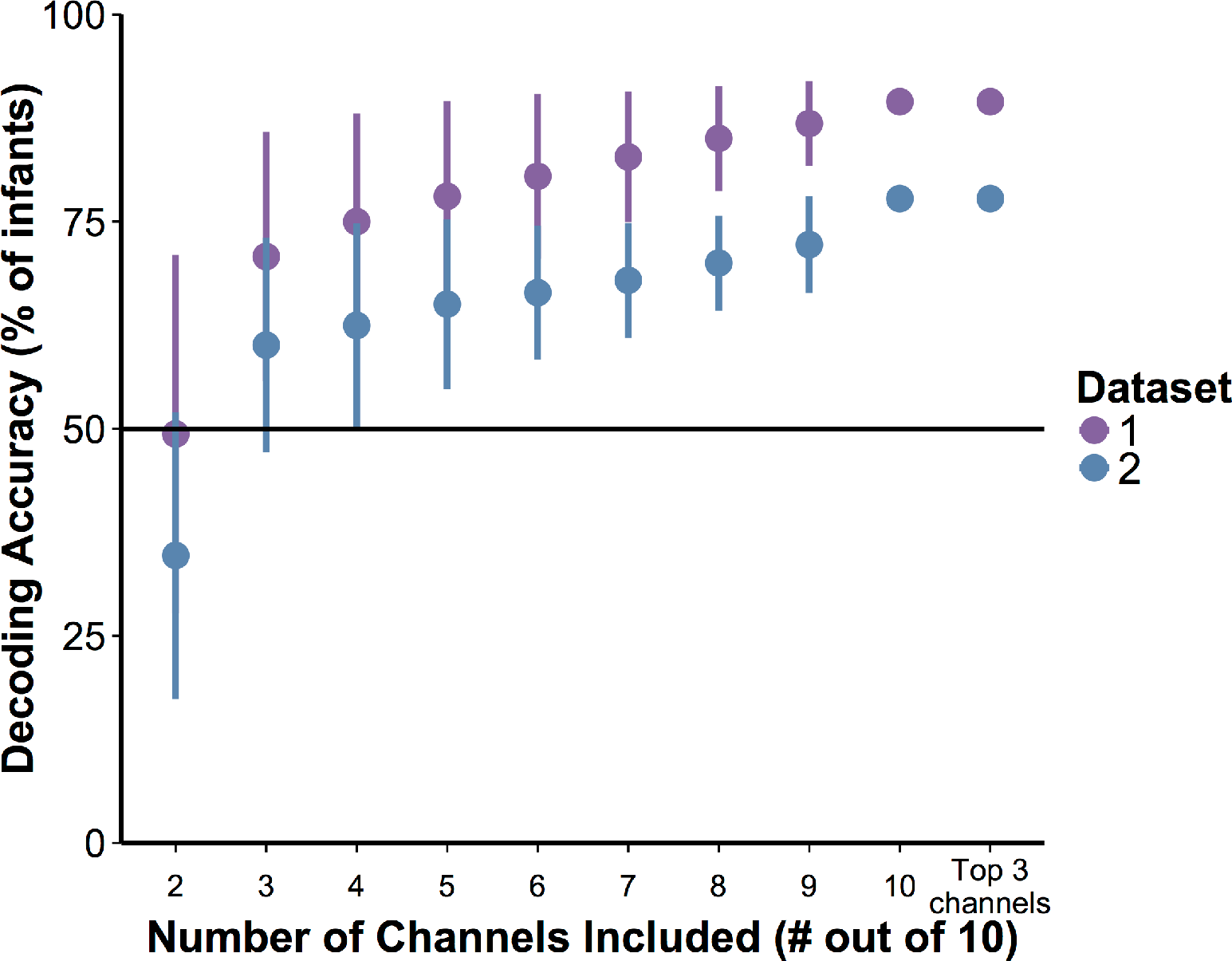
Decoding accuracy of infant-level activation patterns by subset size for Datasets #1 (purple) and #2 (blue). Far right, decoding using three most informative channels (determined using subset size 2, Figure 3, subset size == 3). Error bars are standard deviation of the classification accuracy for all subsets for that size. Note: For subset sizes of 10 channels and Top 3 channels, there is only one subset and so no corresponding standard deviation can be computed. See Supplementary Materials, Figure S3, for a box and whisker version of this plot.

We estimated a logistic regression model to evaluate the following patterns of decoding accuracy for statistical significance:

1. differences in decoding accuracy between Datasets #1 and #2;
2. differences in decoding accuracy across channels;
3. the relationship between subset size and decoding accuracy.

In a mixed-effects logistic regression model (Bates, Maechler, Bolker, & Walker, 2014), we examined infant-level decoding accuracy for each infant, for each subset, and for each channel across subset sizes 2 through 10. In other words, our model attempted to predict decoding accuracy before any of the data had been averaged. Thus, decoding accuracy is either correct, 1, or incorrect, 0, (based on whether the correlation between the infant-level multichannel pattern for the test infant and the group model is higher for the correct labels compared to the incorrect or switched labels, described in detail in Methods). Infants are treated as a random factor in the model to control for any variability across infants. See the Supplementary Material for more details.

First, we confirmed that the difference in accuracy across datasets was significant (coefficient = -0.86, Z = -73.73, *p* < 0.001). Second, by comparing across nested models using chi-squared tests, we confirmed that including channels in our model accounts for significant variance in decoding accuracy (in R mixed-effects model notation: accuracy ~ dataset * channel * set size + (1|Ss), χ^2^(18) = 873.96, *p* < 0.001). This finding indicates that there are systematic decoding differences across channels. We also find that the additional inclusion of subset size in the model significantly increases its fit, indicating that there are differences in decoding accuracy across set size (accuracy ~ dataset * channel * set size + (1|Ss), χ^2^(20) = 2420.6, *p* < 0.001). Third, subset size (from 2 to 10) has a significant positive coefficient in this latter model (coefficient, 0.116, *Z* = 7.95, *p* < 0.001), confirming that with increases in subset size, there is an increase in accuracy (with about a 12% increase in accuracy for each additional channel added, starting with a subset size of 2). These results confirm that there are significant differences in decoding accuracy between datasets, across channels, and with increasing subset size.

### Does the Inclusion of Only the Most Informative Channels Boost Decoding Performance? Functional Evidence for Spatial-Specificity

Reducing the number of channels represented in each multichannel activation pattern decreases decoding accuracy. However, we also found that not all channels are equally informative. In this section, we explore whether the relative informativeness of channels (revealed at smaller subset sizes) translates to better decoding when these channels are combined in a subset. In other words, would the decoding accuracy increase if only the Top 3 most informative channels were used for decoding, compared to what one would expect based on a random sampling of 3 channels? If including only the Top 3 channels boosts decoding accuracy, this would be evidence that channel-by-channel differences in decoding accuracy reflect interregional differences in encoding in the infant brain (i.e., that this method has spatial-specificity).

The most informative channels were defined as the channels with the highest average decoding accuracy at the subset size 2 (see Figure 4; Dataset #1: channels 1, 3, & 8; Dataset #2: channels 1, 3, & 9). Figure 5 graphically presents average decoding performance for these **Top 3 Channels** for the two datasets and indicates nearly identical decoding performance across subjects when compared to using all 10 channels. Moreover, a striking difference can be seen when contrasting the performance of the Top 3 channels to the performance when the subset size is 3 but the average of all such subsets across all channels is computed. To confirm these observations, another mixed effects logistic regression model was estimated to predict decoding accuracy (subset size = 3), when decoding used only the Top 3 most informative channels, and when decoding used all 10 channels (subset size = 10). The statistical model indicated that when controlling for differences across datasets, there were significant differences between the subset size 3 average across all subsets and the Top 3 Channels (*Z* = -2.41, *p* = 0.016) but no differences between the Top 3 Channels and subset size 10 (*Z* = 0.543, *p* = 0.587). See the Supplementary Materials for more details on this model. These results reveal that, while the overall decoding accuracy is reduced at smaller subset sizes, the differences in decoding accuracy across channels at these small subset sizes are significant. When the most informative channels are selected, decoding accuracy is boosted to the level of decoding when all channels are included, and exceed decoding accuracy based on a random selection of channels with the same subset size. These results indicate that MCPA can detect differences in informational processing across regions of the infant brain.

### Comparison to Univariate Analyses: Multivariate analysis distinguishes between conditions when univariate cannot

Multivariate analyses have the ability to exploit the relations between signals from different channels, whereas univariate analyses can only assess each channel individually. Thus, the greatest opportunity for multivariate analyses to reveal information beyond that obtainable by univariate approaches is when the task conditions stimulate broadly distributed regions of the brain at once. In the present study, Dataset #2 contains just such a pair of conditions (faces-and-music, versus fireworks-and-speech). Both conditions are audio-visual, but the specific nature of the audiovisual stimuli differs.

In a univariate analysis of Dataset #2, we compared average activation from 6 to 12.5 seconds after stimulus onset. No single channel exhibited significant differences between conditions after Bonferoni correction (0.05/10 or the number of channels). Thus, there are no robust univariate differences between these two audiovisual conditions. In contrast, our multivariate analysis was able to decode this dataset with high accuracy at the infant-level (> 75%, as depicted in Fig. 3)^5^. This comparison is illustrated in the lower panel of Fig. 6: three channels were jointly able to provide highly accurate multivariate decoding (the three blue-only circles in the Dataset 2 section at the bottom of Figure 6), but there were no channels at all which provided significant univariate decoding (as shown by the absence of any brown rings in that panel). Thus, in trying to distinguish between two types of stimuli, which were both audio-visual but whose content differed (faces-and-music, versus fireworks-and-speech), multivariate analysis was able to distinguish between conditions when univariate could not.

The literature on fMRI decoding has numerous direct comparisons of univariate statistics and multivariate decoding methods (e.g., Kok et al., 2014). At least one previous study of fNIRS decoding has demonstrated the effectiveness of decoding in the absence of significant univariate results (Ichikawa et al., 2014), but that study measured nine-year-old children, a subject group far more cooperative than the infants of our present study, and hence allowed for many more stimulus-presentation trials and much less noisy data than we are using here. Thus, while the benefits of employing MCPA for fNIRS go beyond the benefits of statistical power (see the Introduction), it is important to consider whether MCPA has the potential to uncover neural differences beyond univariate statistics.

**Figure 6.**
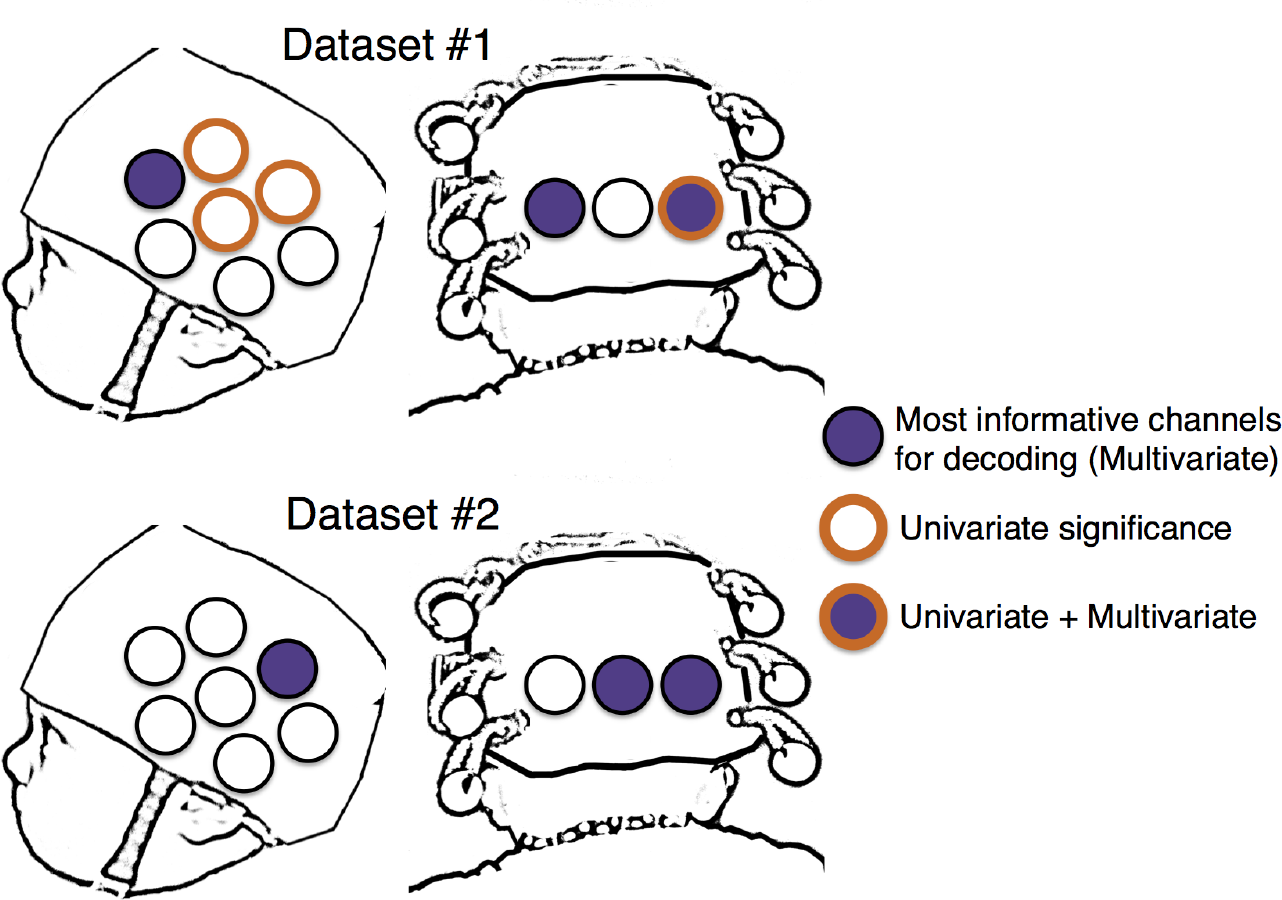
Comparison of the 3 most informative channels for decoding (multivariate “significance”) and channels which exhibit a significant difference between the same two conditions in a univariate analysis. Across both datasets, only a single channel exhibits a significant univariate response and is one of the top channels for the multivariate analyses. In Dataset #2, not a single channel was significant for our univariate analysis.

We also compared univariate and multivariate results in Dataset #1. Average activation was compared between 4 and 9 seconds after stimulus presentation (Emberson et al., 2015). As we expected, there are channels that survive correction in the comparison of unimodal audio and visual stimuli. Specifically, 4 channels exhibited significant differential activity (Channels 1, 7, 9, 10, *ts*(18) > |3.34|, *ps* < 0.0036). Interestingly, these are not the same channels that were the most informative in decoding (i.e., Channels 1, 3 and 8). The channel with the highest relative decoding accuracy (Channel 3) did not exhibit significant differentiation of the two conditions even before statistical correction^6^ Thus, even in the case where infants were presented with unimodal audio and visual stimuli, univariate statistics and MCPA decoding yield different (but highly significant) results, suggesting that these analytic approaches provide different types of information.

## Discussion

We present evidence of significant decoding of the infant brain using a novel multivariate analysis of fNIRS data from two infant datasets. These findings are notable for demonstrating a simple, effective multivariate method for fNIRS data with highly accurate decoding of neural responses across infants that either complements or surpasses univariate analyses. Specifically we proposed a correlation-based analytic framework for conducting a multivariate analysis on a small set (n=10) of NIRS channels. The resultant multichannel pattern analysis (MCPA) reliably decoded which of two stimulus conditions was present (i.e., the average pattern of response across channels) both at the infant-level and at the trial-level (i.e., the average pattern for an infant across all trials and the average pattern for a single trial, respectively). Notably, decoding was conducted between-infants (i.e., based on a group model that did not include data from the test infant).

Infant-level performance was robust across two datasets, which included both unimodal and multimodal stimuli. Trial-level performance was reliable for unimodal stimuli, and despite not achieving statistical significance, trial-level prediction accuracy in the multimodal stimuli (60%) was strongly suggestive of subtle differences in the sensory properties of the two stimulus conditions. Future trial-level discrimination may simply require more statistical power (i.e., larger sample size of infants). Nevertheless, decoding accuracy using MCPA was highly robust under for decoding across infants in both unimodal and multimodal stimulus conditions. This result is especially impressive given the many hurdles that must be overcome when gathering fNIRS data from infants (e.g., limited number of channels, large spatial extent of sampled cortex per channel, small number of trials contributed per infant, greater intersubject variability).

The subset analyses suggest that MCPA methods benefit from including a greater numbers of channels, highlighting the multivariate aspect of this method. Further, this analysis revealed **differential informativeness of individual channels** to MCPA decoding accuracy such that—after identifying the most informative channels—the same level of decoding accuracy could be achieved with a select subset of informative channels. This finding in particular establishes that spatially-specific information about representations in the brain can be achieved using MCPA. Moreover, both of these findings validate the multivariate method as a tool for identifying pattern-based neural correlates of cognition: The decrease in average decoding accuracy with smaller subset sizes establishes that the current analysis method utilizes patterns of activation distributed across channels and thus suffers when fewer of these channels are contributing to the observed pattern of activation. It also follows that not all channels contribute equally to decoding, and thus channel-wise analyses indicate differences in specialization or engagement of specific cortical regions to the current task, which is detectable using MCPA.

The most informative channels identified in the present study were widely distributed across the cortical surface including occipital, temporal and frontal regions. This distribution of informative channels highlights the value of MCPA for pooling information across the brain without spatial constraint. The evaluation of subset informativeness opens a window on exploring the continuum of modular (i.e., cluster-based) versus distributed neural architectures, including how such architectures might be biased before birth and/or emerge with exposure to postnatal experience. This inference about spatial localization intersects with the goals of univariate analyses, which focus on clusters of channels (i.e., an ROI approach). However, a direct comparison of univariate and multivariate methods in this study revealed that MCPA elucidates aspects of the hemodynamic signal that are missed in standard univariate tests. In the multimodal dataset (Dataset #2), significant infant-level decoding is successful using a subset of informative channels while univariate methods fail to find any significant differences on these (or indeed, any) channels. Moreover, in both datasets, the individual channels that were demonstrably the most informative in decoding did not strongly overlap with the channels that exhibited significant univariate results. In fact, only a single channel both survived statistical correction in univariate tests and was one of the top 3 channels for decoding performance (Channel 3 in Dataset #1). Thus, on two fronts we find that MCPA notably extends results found using univariate methods: In the absence of significant univariate results, decoding is still possible (Dataset #2) and individual channels can be identified as supporting decoding while yielding no significant univariate results and vice versa.

Given this initial proof of concept that MCPA can be used successfully to decode two stimulus conditions from the fNIRS data of 6-month-old infants and provide findings beyond univariate tests, there are a large number of questions that can be tackled which were hitherto inaccessible to fNIRS studies using univariate methods. For example, rather than asking whether cortical region X is more activated by a particular type of stimulus (e.g., faces) than some other comparison stimulus (e.g., houses), one can ask two more subtle questions. First, is there a *pattern* of activation that, even in the absence of a difference in mean activation for the two stimulus conditions, indicates reliable decoding of that stimulus difference? Second, given a reliable pattern of decoding, what is the spatial distribution of fNIRS channels that are *informative* for that decoding process? Answers to these two questions, which our findings now bring into the realm of possibility, will enable researchers who study the *development* of the human brain to begin to propose and evaluate more sophisticated models of the neural mechanisms of cognition. In addition, by relating the patterns of fNIRS activations among stimuli that vary along known dimensions, one can expand MCPA to ask how higher-level stimulus dimensions are decoded by the brain. Now that MCPA has been confirmed as a viable method in infants, future use of Representational Similarity Analysis promises to be a fruitful avenue for investigation, such as implementing methods for more than two conditions, as described by Raizada & Connolly (2012) and Anderson, Zinszer, & Raizada (2015).

Aslin et al.'s (2015) review of fNIRS contribution to developmental research highlighted the imminent need to extend multivariate methods developed in fMRI to fNIRS. Although some previous studies have implemented machine learning techniques to classify fNIRS responses, these examples have not yet been widely adopted by the fNIRS community, and we have argued that there are significant barriers to doing so both computationally and interpretatively. In this paper and the supplementary material, we offer a simple, transparent method for decoding fNIRS data in an early developmental population. The MATLAB code for implementing these analyses is publicly available, flexible enough to easily accommodate various experimental configurations (e.g., different numbers of fNIRS channels or subsets of channels), and can automatically extract analysis-relevant details from HomER data files (Huppert, Diamond, Franceschini, & Boas, 2009). The availability and transparency of the methods and code will also allow researchers to make changes that expand the use of MCPA at any stage of the analysis pipeline for multivariate fNIRS research.

## Footnotes

1 http://teammcpa.github.io/EmbersonZinszerMCPA/

2 Ichikawa et al. (2014) overcame this limitation to decode clinical diagnoses (ADHD/autism) in children with a classifier that required a large amount of computational power. The training necessary for this technique required the calculation of 2^24^ or 17 million subsets of the data.

3 As we are decoding over averaged windows of oxygenated hemoglobin, in general, there is not yet a clear and principled way to select time-windows for the current analyses. Future work could identify optimal timing for event designs (as in Dataset 1) versus block designs (as in Dataset 2) or model beta values as in adult fMRI analyses. The application of multivariate methods to infant fNIRS data is indeed in its infancy, and the current study leaves these avenues to further research.

4 It is also interesting to note that for Dataset #1 while the differences between channels becomes more prominent as subset size decreases, the overall pattern of which channels are more informative compared to others remains stable. This is not the case for Dataset #2: Between subset sizes of 2 and 3, there are differences in which channels are (relatively) more or less informative particularly in which channels are the least informative.

6 An additional 3 channels exhibited significant differences (p < 0.05) but did not survive correction (Channels 4, 5 and 8).

## Acknowledgements

The authors wish to thank all the caregivers and infants who volunteered their time to make this research possible as well as Holly Palmeri, Ashley Rizzieri, Eric Partridge, Kelsey Spear and the rest of the research assistants in the Rochester Baby Lab. This work was supported by an NICHD Grant K99 HD076166-01A1, 4R00HD076166-02, and a Canadian Institutes of Health Research (CIHR) postdoctoral fellowship to (to L.L.E.), NSF Award 1228261 (to R.R) and NIH Grant R01 HD-37082 and NSF EAGER BCS-1514351 (to R.N.A.).

## Supplementary Materials

### NIRS Channel Locations: MR-coregistration

Across both datasets, the same NIRS cap configuration was employed. MR co-registration for this cap configuration has been conducted for over 100 infants to reveal the average cortical localization for each NIRS channel. Table 1, adopted from Emberson et al., (2015), presents the localization for a representative subset of these infants and including infants in Dataset #1. See Emberson et al., (2015) for detailed descriptions of the MR-coregistration methods employed.

It is important to note that while the MR-coregistration methods employed provide evidence of the average spatial location, there are certainly differences across individual infants in the precise cortical regions being recorded from a given channel for a number of reasons. First, the NIRS cap is a fixed size and configuration so, despite efforts to standardize placement of the cap for each infant, any change in head size will result in differences in the cortical regions being recorded for each channel. Second, individual differences in neuroanatomy are not accessible to fNIRS because unlike fMRI no anatomical image of each brain is able to be collected. Given that MCPA employs between-subjects decoding, it is a strong assumption of the method that the same channel is recording from the same cortical regions across the dataset. However, given the large size of each fNIRS channel (~2cm of cortex), it is not clear how much these variations matter for MCPA decoding or, indeed, any fNIRS data analysis. Of course, better spatial specificity in the recordings will allow MCPA to reveal more precise spatial specificity in the dataset, and if a researcher makes no attempt to control or at least quantify the locations of the fNIRS channels, this will certainly reduce the accuracy of decoding. However, we believe that the major constraint on spatial specificity for MCPA is the size of the fNIRS channels and the lack of collection of individual neuroanatomical images for each subject.

**Table 1:** Localization of NIRS Channels.

| Channel No. | MR Type | Lobar Atlas (%) | LONI Atlas (%) |
| --- | --- | --- | --- |
| **Channel-1** | average | Occipital (79) | Middle occipital gyrus (49) |
|  | individual | Parietal (88) | Middle occipital gyrus (52) |
| **Channel-2** | average | Occipital (88) | Middle occipital gyrus (37) |
|  | individual | Parietal (52) | Middle occipital gyrus (58) |
| **Channel-3** | average | Parietal (47) | Middle occipital gyrus (63) |
|  | individual | Parietal (79) | Middle occipital gyrus (73) |
| **Channel-4** | average | Temporal (92) | Middle temporal gyrus (48) |
|  | individual | Temporal (97) | Middle temporal gyrus (55) |
| **Channel-5** | average | Temporal (90) | Middle temporal gyrus (44) |
|  | individual | Temporal (97) | Middle temporal gyrus (58) |
| **Channel-6** | average | Temporal (86) | Superior temporal gyrus (43) |
|  | individual | Temporal (94) | Superior temporal gyrus (64) |
| **Channel-7** | average | Temporal (65) | Middle temporal gyrus (41) |
|  | individual | Temporal (67) | Middle temporal gyrus (61) |
| **Channel-8** | average | Frontal (67) | Inferior frontal gyrus (34) |
|  | individual | Frontal (88) | Precentral gyrus (52) |
| **Channel-9** | average | Temporal (51) | Superior temporal gyrus (30) |
|  | individual | Parietal (52) | Supramarginal gyrus (47) |
| **Channel-10** | average | Frontal (43) | Postcentral gyrus (34) |
|  | individual | Parietal (58) | Postcentral gyrus (47) |

### Permutation-Based Significance Tests

The assumptions for the binomial test are not satisfied in *n*-fold cross validation (i.e., leave-one-infant-out cross validation for all *n* infants) due to the non-independence of each fold (violates the *i.i.d.* assumption of a parametric test). Thus it is necessary for us to generate an empirical null distribution composed of either all the possible outcomes under the null hypothesis or a large, random sample of those outcomes. This null distribution is estimated by permuting the labels on each subject’s activation patterns for the two conditions and computing the distribution of decoding accuracies across many (e.g., 10,000) such permutations (see Nichols & Holmes, 2003 for background).

#### Infant-Level Decoding

Because there are two conditions, the number of possible label permutations in this dataset is 2^*n*^ where *n* is the number of subjects. Because the infant-level decoding is computationally fast (<1 sec to complete the entire *n*-fold cross validation), we were able to explore all possible permutations for each dataset (2^7^ for Dataset #1 and 2^18^ for Dataset #2). In this case, the null distributions represent *all* possible label permutations, and the null hypothesis is that infant-level decoding accuracy with the correct condition labels does not significantly differ from randomly assigned labels.

Figure S1 depicts frequencies (histogram divided by total number of permutations) for the accuracy of the infant-level permutation tests of Dataset #1 (left panel) and Dataset #2 (right panel). P-values for the observed decoding accuracies (Dataset #1: 17/19 infants correctly labeled; Dataset #2: 14/18 infants correctly labeled) are computed by measuring what proportion of observations are equal to or greater than the observed decoding accuracy on the null distribution. For reference, all values on the distribution with *p*<0.05 are highlighted in red.

**Figure S1.**
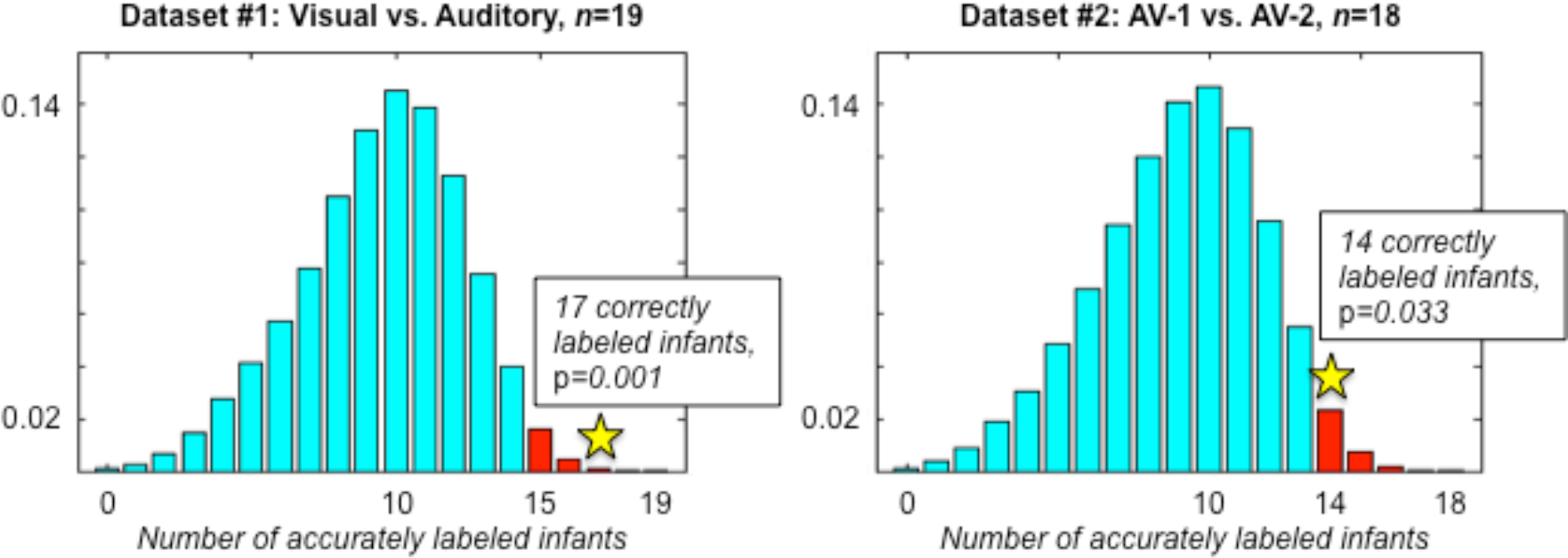
Null distributions for infant-level decoding in Dataset #1 and Dataset #2. Values with empirical *p*-value less than 0.05 are highlighted in red, and observed accuracies reported in Results section are indicated with a star and labeled.

#### Trial-Level Decoding

The permutation procedure for trial-level decoding was the same as infant-level decoding,but instead of measuring the number of correctly labeled infants, the number of correctly labeled trials for each condition was averaged across infants in the cross validation (following the same method as computing actual accuracy described in the Methods). Because trial-level decoding requires greater computation time (approximately 2 seconds per *n* folds), searching the entire set of possible permutations was computationally infeasible. Instead, we randomly sampled 10,000 permutations from the set of all possible permutations. Mean trial-level accuracy across infants is calculated for each condition in each permutation. The null distribution thus represents the average trial-level decoding accuracy when labels are randomly assigned.

Figure S2 depicts the null distributions for each of the two conditions in Datasets #1 (top two panels) and #2 (bottom two panels). *P*-values for the observed decoding accuracies (Dataset #1: Condition 1 68%, Condition 2 66%; Dataset #2: Condition 1 60%, Condition 2 57%) are again computed by measuring what proportion of observations are equal to or greater than the observed decoding accuracy on the null distribution. For reference, all values on the distribution with p<0.05 are highlighted in red.

**Figure S2.**
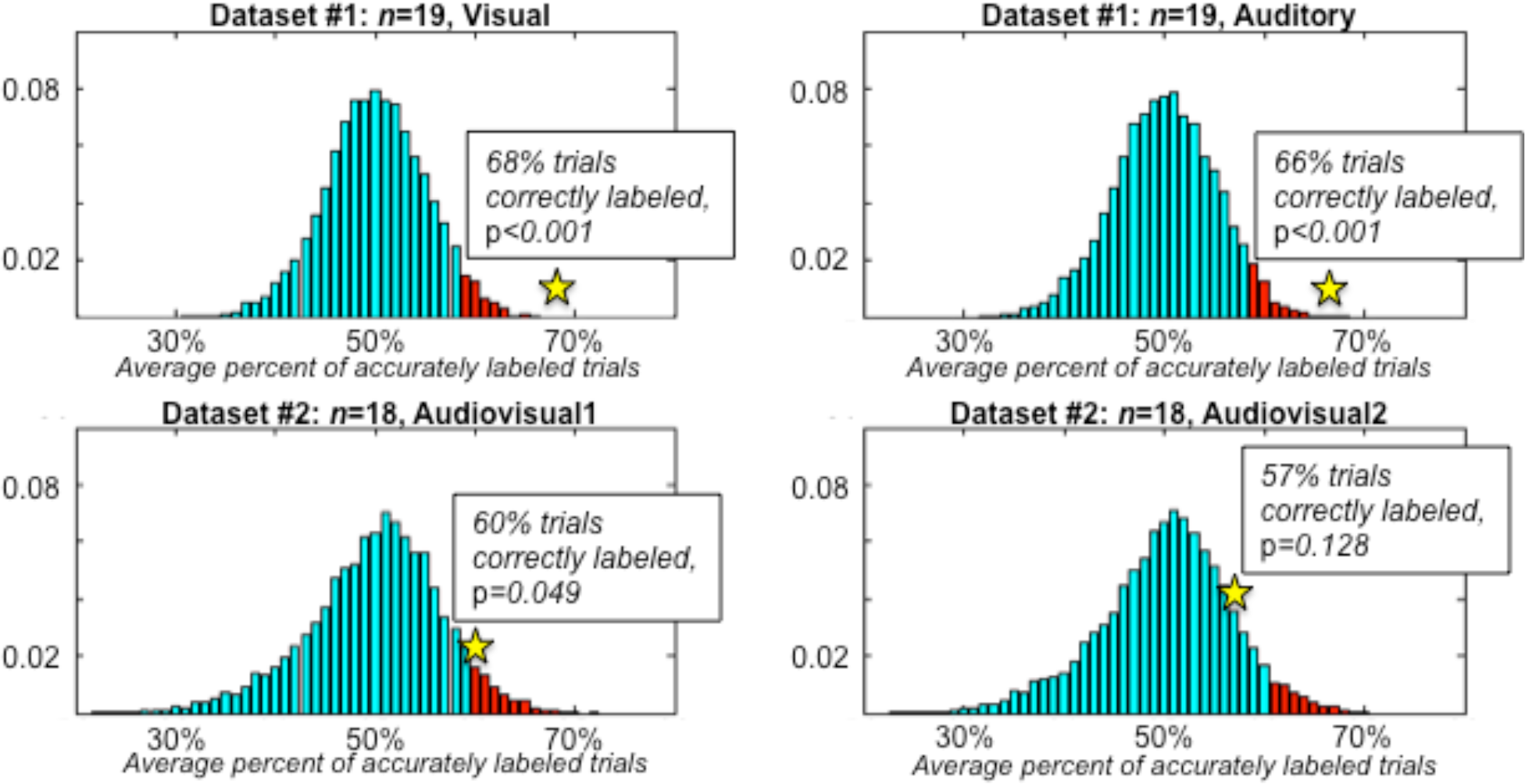
Null distributions for trial-level decoding for each condition in Dataset #1 and Dataset #2. Values with *p*<0.05 are highlighted in red, and observed accuracies reported in Results section are indicated with a star and labeled.

### Models of Decoding Accuracy

In subsection “Are there differences in decoding accuracy with subset size?”, the data modeled is the decoding accuracy (1/0) for each infant, for each subset, for each channel, for each dataset which allows us to determine whether there are differences in decoding accuracy across dataset, across channels, and what the relationship is between subset size and decoding accuracy. The employed the glmer function in the lme4 package in R (Bates et al., 2014). Here we include pseudo-code to illustrate the details of the model. We've made both this data and the code available for more detailed analysis, replication, and extension. Results and coefficients of these models are reported in the main text of the paper.

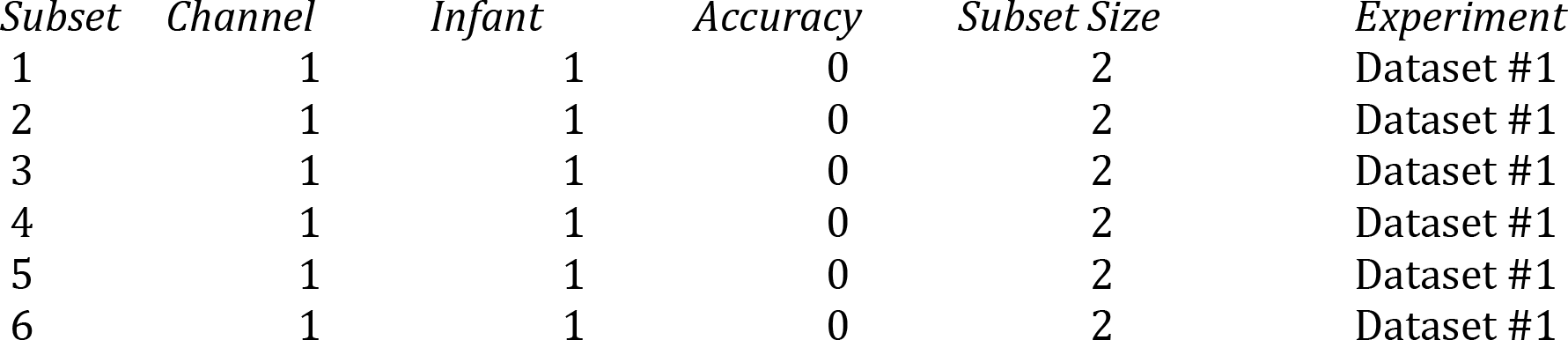

| *Subset* | *Channel* | *Infant* | *Accuracy* | *Subset Size* | *Experiment* |
| --- | --- | --- | --- | --- | --- |
| 1 | 1 | 1 | 0 | 2 | Dataset #1 |
| 2 | 1 | 1 | 0 | 2 | Dataset #1 |
| 3 | 1 | 1 | 0 | 2 | Dataset #1 |
| 4 | 1 | 1 | 0 | 2 | Dataset #1 |
| 5 | 1 | 1 | 0 | 2 | Dataset #1 |
| 6 | 1 | 1 | 0 | 2 | Dataset #1 |

We start by determining whether there are differences in decoding accuracy across <u>datasets</u> (with Dataset #1 set as the intercept).

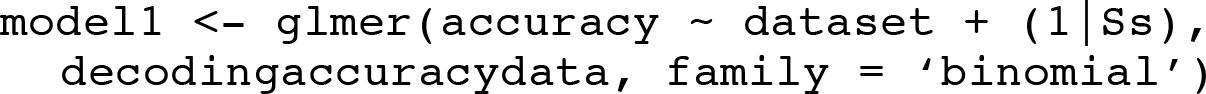

When then add the additional factor of <u>channel</u> to the model (channel #1 as the intercept) and evaluate whether this addition significantly predicts more of the variance as a significant increase in model fit would indicate that decoding accuracy is not equal across channels.

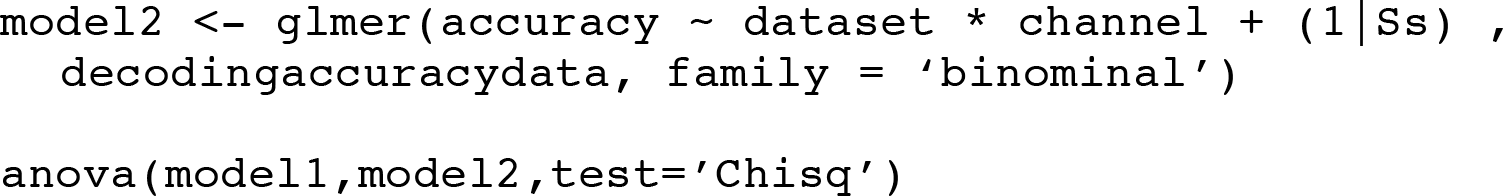

We then determine whether the addition of <u>subset size</u> (numeric predictor) significantly improves the model and if so what kind of relationship <u>subset size</u> has on decoding accuracy (e.g., is it a significant positive relationship as indicated graphically).

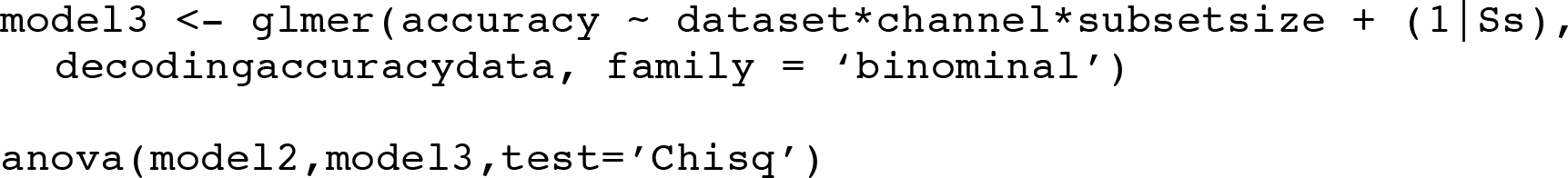

In subsection “Does the inclusion of only the most informative channels boost decoding performance?” we again model the decoding accuracy for each subset, channel and infant but only for subset sizes 3, 10 and Top 3 Channels. The inclusion of decoding accuracy for the Top 3 Channels and the restriction of subset sizes to 3 and 10 allows a more direct comparison of decoding accuracy. In these models, both infant and channels are treated as random factors as variation in these factors is not of experimental interest and not including them in the model could result in inflated p values.

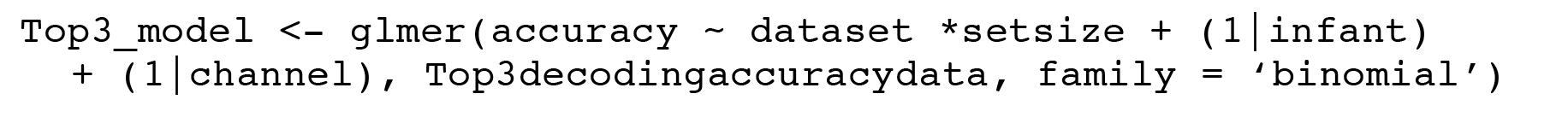

See github.com/laurenemberson/EmbersonZinszerMCPA_analysesFromPaper for code to implement these models as well as the decoding accuracies from the two datasets reported in this paper.

**Figure S3.**
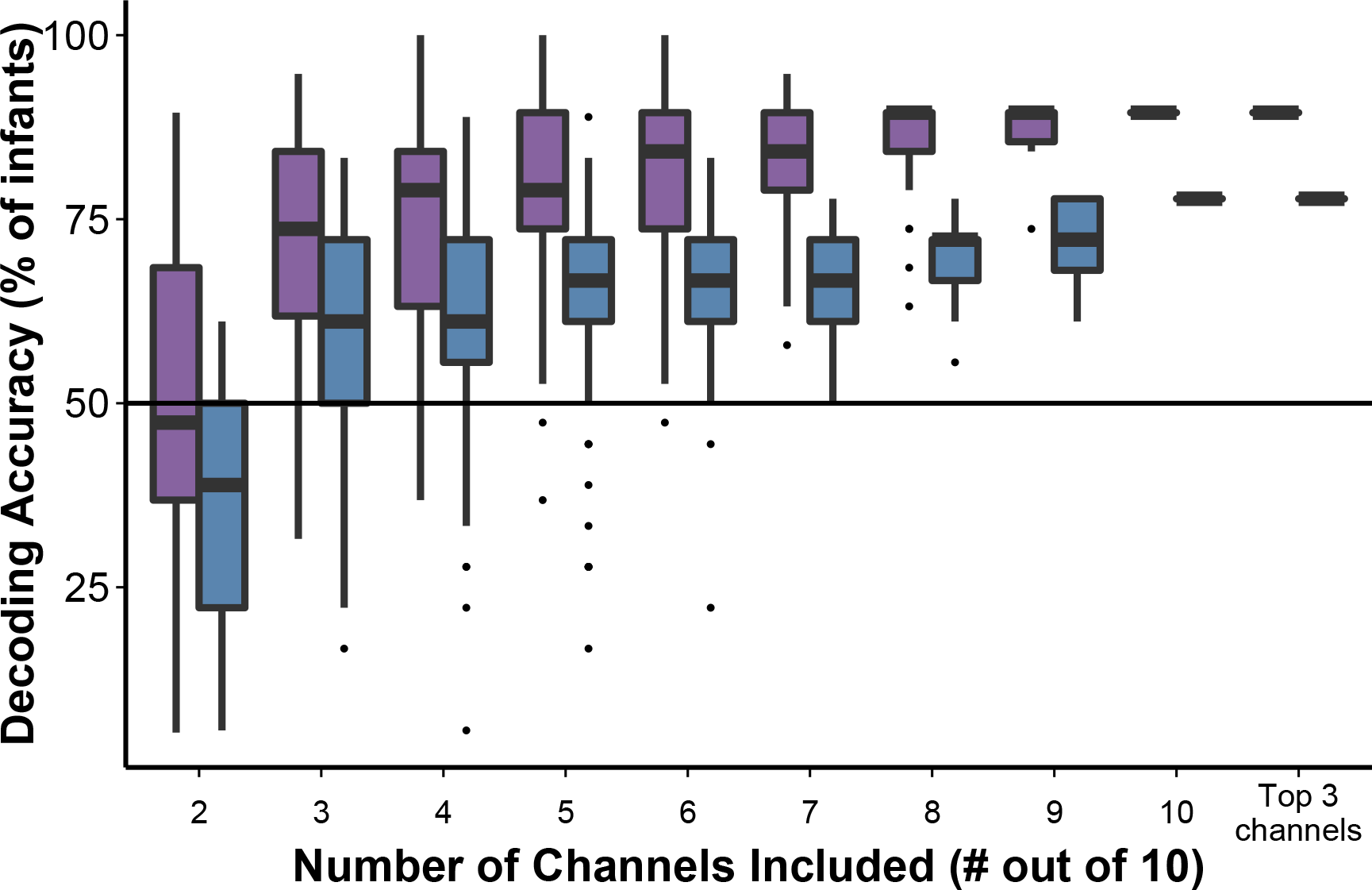
Decoding accuracy of infant-level activation patterns by subset size for Datasets #1 (purple boxes) and #2 (blue boxes). Far right, decoding using three most informative channels (most informative channels determined using subset size 2, Figure 3). Note: For subset sizes of 10 channels and Top 3 channels, there is only one subset and so there is no range to estimate. Figure corresponds to Figure 4 in the main text.

